# Deep-SMOLM: Deep Learning Resolves the 3D Orientations and 2D Positions of Overlapping Single Molecules with Optimal Nanoscale Resolution

**DOI:** 10.1101/2022.07.31.502237

**Authors:** Tingting Wu, Peng Lu, Md Ashequr Rahman, Xiao Li, Matthew D. Lew

## Abstract

Dipole-spread function (DSF) engineering reshapes the images of a microscope to maximize the sensitivity of measuring the 3D orientations of dipole-like emitters. However, severe Poisson shot noise, overlapping images, and simultaneously fitting high-dimensional information–both orientation and position–greatly complicates image analysis in single-molecule orientation-localization microscopy (SMOLM). Here, we report a deep-learning based estimator, termed Deep-SMOLM, that archives superior 3D orientation and 2D position measurement precision within 3% of the theoretical limit (3.8^◦^ orientation, 0.32 sr wobble angle, and 8.5 nm lateral position using 1000 detected photons). Deep-SMOLM also achieves state-of-art estimation performance on overlapping images of emitters, e.g., a 0.95 Jaccard index for emitters separated by 139 nm, corresponding to a 43% image overlap. Deep-SMOLM accurately and precisely reconstructs 5D information of both simulated biological fibers and experimental amyloid fibrils from images containing highly overlapped DSFs, at a speed ∼10 times faster than iterative estimators.

## 1. Introduction

Single-molecule orientation-localization microscopy (SMOLM) is a versatile tool for visualizing interactions between biomolecules and the architecture of the structures they create; it measures simultaneously the 3D orientations and positions of individual fluorescent molecules with nanoscale resolution. Researchers have used molecular orientations [1] to elucidate the architectures of amyloid aggregates [2–4], the organization of proteins in actin filaments [5, 6], and changes in lipid membrane polarity and fluidity induced by cholesterol [4, 7, 8]. To effectively use limited photon budgets in single-molecule (SM) imaging, dipole-spread functions (DSFs), i.e., the vectorial extension of the optical microscope’s point-spread function, must be engineered to encode additional information about a molecule’s 3D orientation [5, 9–15]. However, simultaneously estimating the 3D orientation and position of an emitter is challenging because 1) it is difficult to estimate SM parameters in 5-dimensional space (3D orientation, wobble, 2D position) without getting trapped in local minima instigated by severe Poisson shot noise [4, 14], 2) engineered DSFs have larger footprints and cause SM images to overlap frequently, and 3) dim emitters are difficult to detect when large DSFs spread photons across many camera pixels.

To estimate SM orientations, existing techniques match noisy experiment images to a precomputed library of sampled DSFs [16] or construct parametric fits using models of the imaging system [5, 9–11, 13, 17, 18]. These methods either suffer from 1) reduced precision due to finite sampling and/or DSF approximation or 2) a heavy computational burden during the slow, iterative optimization process. In addition, estimating parameters of dim SMs whose images overlap is extremely challenging. Fundamentally, estimating 3D orientation and 2D position is equivalent to deconvolving five-dimensional features from two-dimensional noisy images. Convolutional neural networks have achieved great success in shift-invariant pattern recognition [19–21]. Early neural networks for measuring orientation are limited to images containing one emitter [22, 23]. Recently, DeepSTORM3D and DECODE have been developed for estimating the 3D positions of single molecules, even for a high density of emitters whose images overlap [24, 25]. However, techniques that are capable of high-dimensional estimates, i.e., measuring five or more parameters, from overlapping images of dense emitters are still missing.

In this paper, we propose a deep-learning based estimator, termed Deep-SMOLM, for simultaneously estimating the 3D orientations and 2D positions of single molecules. Through carefully designed synergy between the physical forward imaging model of the microscope and the neural network architecture, Deep-SMOLM estimates both SM positions and orientations on par with the theoretical best-possible precision at a speed ∼10 times faster than iterative estimators. Morever, Deep-SMOLM accurately and precisely analyzes overlapping images of dense emitters in both realistic imaging simulations and in experimental imaging of amyloid fibrils. To the best of our knowledge, Deep-SMOLM is the first deep-learning based estimator capable of estimating 5D information from overlapping images of single molecules.

## 2. Methods

We represent the mean orientation of a dipole-like emitter [26–28] using a polar angle *θ* and an azimuthal angle *ϕ* in spherical orientation space (Fig. 1(a)). During a camera exposure, the dipole rotates or “wobbles” through a range of directions represented by a solid angle Ω (Fig. 1(a)) [29,30]. Traditional unpolarized microscope images contain little information about the 3D orientation of a dipole [1, 12, 31, 32]. To remedy these issues, we use a polarization-sensitive microscope that splits fluorescence into x- and y-polarized detection channels, along with a pixOL phase mask placed at the pupil or back focal plane (Fig. 11) [14]. The resulting pixOL DSF changes dramatically for molecules oriented in various directions (Fig. 1(b)).

**Fig. 1.**
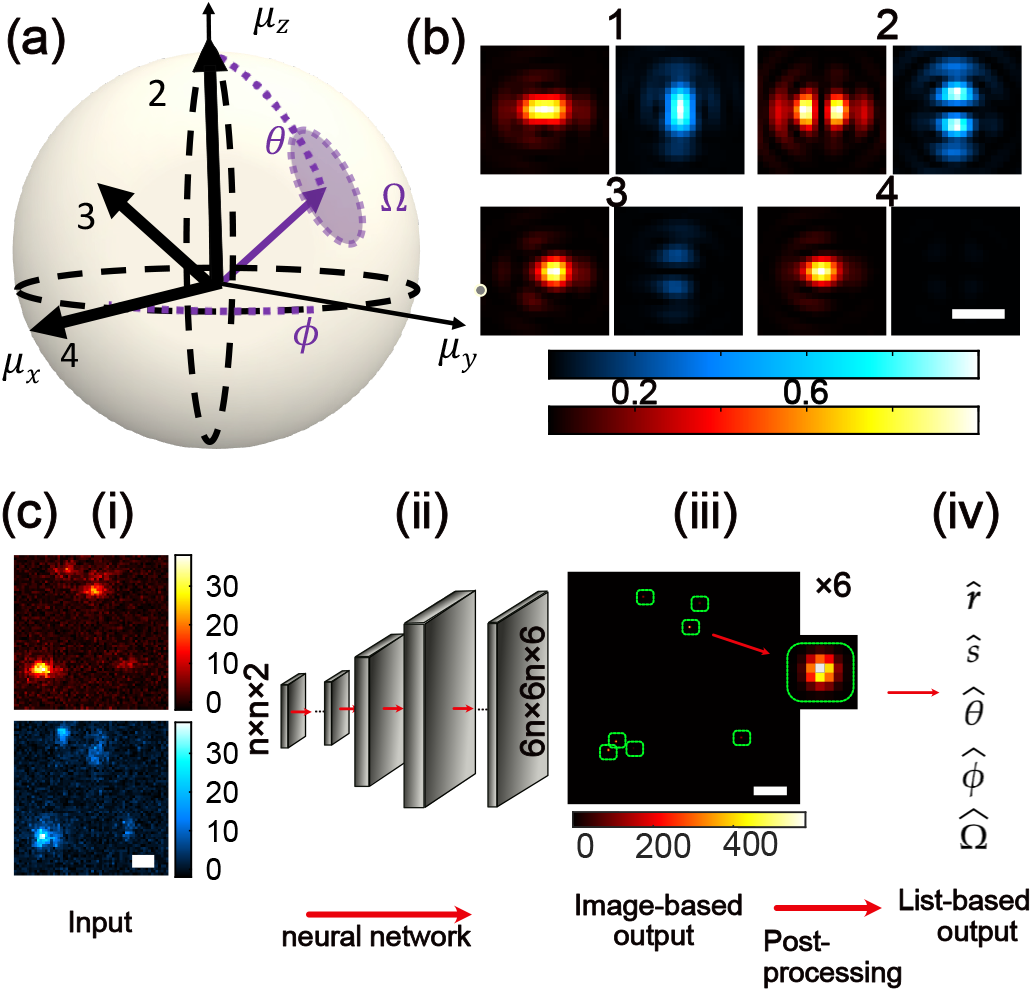
Estimating 3D orientations and 2D positions of single molecules (SMs) using Deep-SMOLM. (a) The orientation of a dipole-like emitter is parameterized by a polar angle *θ* ∈ [0, 90^◦^], an azimuthal angle *ϕ* ∈ (−180^◦^, 180^◦^], and a wobble solid angle Ω ∈ [0, 2*π*] sr. (b) Simulated (left, red) x- and (right, blue) ypolarized images captured by a polarization-sensitive microscope with a pixOL phase mask [14] of emitters with orientations [*θ, ϕ*, Ω] shown in (a). Emitter 1: Ω = 2*π* sr; emitter 2: [0^◦^, 0^◦^, 0]; emitter 3: [45^◦^, 0^◦^, 0 sr]; emitter 4: [90^◦^, 0^◦^, 0 sr]. (c) Schematic of Deep-SMOLM. (i) A set of (top, red) x- and (bottom, blue) y-polarized images of size of *n* × *n* is input to (ii) the neural network, which outputs (iii) six images ***h*** of size 6*n* × 6*n* (Eqn. 3). Each detected emitter is represented by a 2D Gaussian spot ((iii) inset) located at corresponding positions across the six images. An emitter’s 2D position 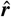 is encoded into the center position of the Gaussian pattern, and the signal-weighted moments are encoded as the intensities of the Gaussian patterns across the six images ***h***. (iv) A post-processing algorithm is used to transform the Deep-SMOLM images into a list of SMs, each with a measured 2D position 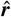, intensity *ŝ*, and 3D orientation 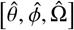.

To estimate these orientations, the network must detect a DSF above background in the presence of Poisson shot noise and estimate the 3D orientation and 2D position of a molecule based on the shape of the DSF. Directly estimating orientation angles is generally ill-conditioned: the mean orientation direction [*θ, ϕ*] is periodic [33, 34] and occasionally degenerate, e.g., for an isotropic emitter (Ω = 2*π*), the mean orientation angle [*θ, ϕ*] is undefined. To mitigate these issues, Deep-SMOLM estimates the brightness-weighted orientational second moments 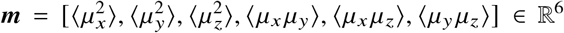 instead of orientation angles (see SI Section 1 for the relation between orientation angles and second moments). The image of an SM produced by the microscope is linear in these second moments, as modeled by vectorial wave diffraction [35, 36], such that

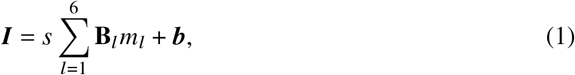

where ***I*** ∈ ℝ^*N*×*P*^ is the measured fluorescence intensity within a pair of x- and y-polarized images with *N* × *P* pixels, *s* is the number of signal photons detected from the emitter, and ***b*** is the background in each pixel. The *l*^th^ basis image **B**_*l*_ ∈ ℝ^*N*×*P*^ correspond to the imaging system’s response to *l*^th^ orientational second moment *m*_*l*_ (Fig. S7). Importantly, while the image ***I*** of an SM is linear with respect to the second moments ***m***, the second moments are nonlinear with respect to the orientation angles (SI Section 1 or Eqn. S1). Moreover, there exists a unique one-to-one mapping between orientational second moments ***m*** and SM images ***I*** (Eqn. 1), but a single image can be produced by multiple orientation angles [*θ, ϕ*, Ω].

Extending the forward model (Eqn. 1) to images containing *Q* emitters with the *q*^th^ emitter located at ***r***_*q*_ = [*x*_*q*_, *y*_*q*_], we have

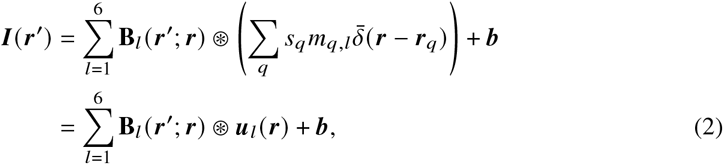

where 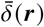 is the 2D Dirac delta function, *m*_*q,l*_ is the *l*^th^ orientational second moment for *q*^th^ emitter, and ⊛ is the convolution operator. Thus, both the 3D orientations and 2D positions of all emitters within a camera frame can be explicitly and uniquely represented by ***u***(***r***), a six-dimensional vector field defined for all possible molecule positions ***r***. Each entry of ***u***_*l*_ (***r***) corresponds to one brightness-weighted orientational second moment. Deep-SMOLM specifically leverages the linearity of the imaging model in Eqn. 2 to estimate brightness-weighted orientational second moment images ***u***(***r***) for improved estimation performance.

It is impossible for a network to estimate ***u***(***r***) containing Dirac delta functions within a continuous 2D domain ***r***. Inspired by Deep-STORM and Deep-STORM3D [24,37], we designed Deep-SMOLM to instead generate six Gaussian-blurred images ***h***_*l*_ (***r***), each corresponding to a brightness-weighted second moment image, as

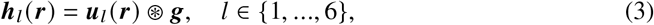

where ***g*** is a Gaussian kernel with a width (standard deviation) of 1 pixel represented by a 7 × 7 matrix. The spacing of pixel grid is 9.7 nm, close to the localization precision of single molecule imaging. Thus, Deep-SMOLM represents each detected emitter by 2D Gaussian spots co-located within each of its 6 output images ***h***_*l*_ (***r***) (Fig. 1(c)(iii) and Fig. S3). To compile a list of position and orientation estimates, each corresponding to a detected SM, a post-processing algorithm simply identifies and crops each Gaussian pattern. The SM’s 2D position is computed from the center of the Gaussian pattern, and the SM’s orientation and intensity are measured from the amplitudes of the Gaussian patterns across the six images ***h***_*l*_ (***r***) (Fig. 1(c)(iv), see SI Section 2.iii for more details).

We train the network using 30K simulated images, each containing a random number of molecules, with a density between 0.6 and 1.2 molecules·*µ*m^−2^, placed at random positions with random orientations (SI section 4.ii). The images contain both well-separated and overlapped DSFs. The network is optimized by minimizing the loss given by

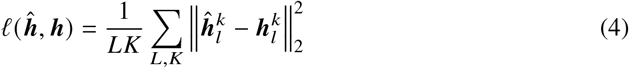

where 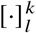 represents the *k*^th^ pixel of the *l*^th^ ground truth image ***h, ĥ*** is the output images from the network, and 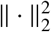 is the l2 norm. The same loss function is used throughout all training epochs for testing dataset to evaluate the performance. Full network training takes ∼2 h on a Nvidia GeForce RTX 2080 Ti GPU with 11 GB memory.

## 3. Results

### i. Estimation accuracy and precision

In contrast to image-based algorithms, which output discrete position estimates and are therefore limited to pixel-level precision [24, 37, 38], Deep-SMOLM encodes the 2D position of an SM into the center of a Gaussian spot in its output images, and a post-processing algorithm outputs a continuous 2D position estimate ***r*** = [*x, y*]. To test localization accuracy, we place four emitters at various locations with respect to the output pixel grid (spacing = 9.7 nm), namely at 0 nm, 2.9 nm, 5.9 nm, and 8.8 nm. For each emitter, we generate 2000 noisy images, each containing one emitter with a random orientation sampled from a uniform distribution and with 1000 detected signal photons and 2 background photons/pixel. The estimated positions from Deep-SMOLM for the four emitters are centered at their ground-truth positions (Fig. 2(a)) with an average bias *x*_bias_ of -0.14 nm.

**Fig. 2.**
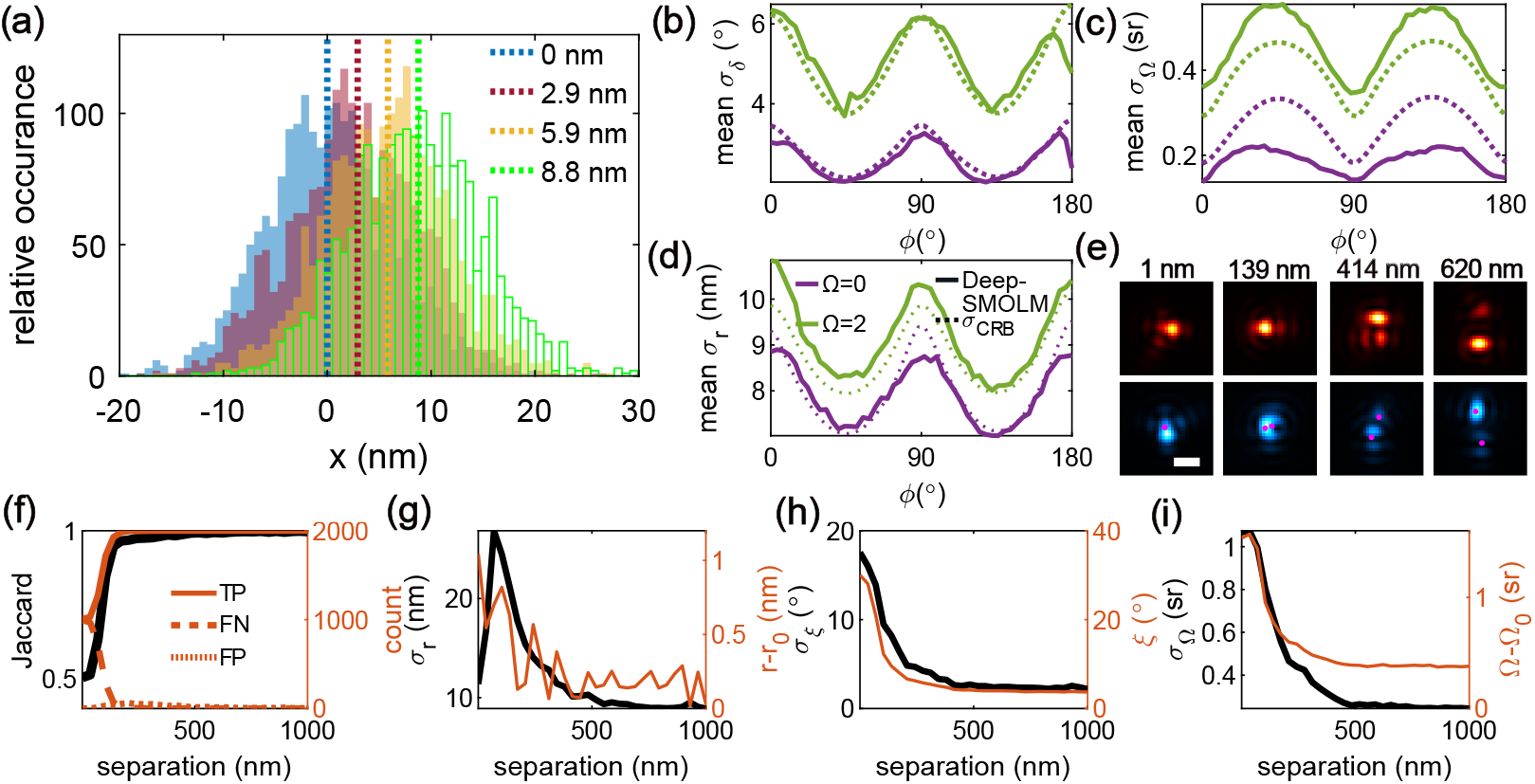
Precision of Deep-SMOLM for estimating 3D orientations and 2D positions of SMs. (a) Estimated lateral position *x* for emitters located at (blue) *x* = *y* = 0 nm, (red) 2.9 nm, (yellow) 5.9 nm, and (green) 8.8 nm. For each case, 2000 noisy images are simulated containing an SM located at the designed position with a random orientation. Deep-SMOLM measurement performance, as quantified by mean angular standard deviation *σ*_*δ*_ (MASD, Eqn. S7), wobble angle precision *σ*_Ω_, and lateral precision *σ*_*r*_ averaged uniformly over all *θ*. Solid line: Deep-SMOLM precision, dashed line: Cramér-Rao bound precision, purple: Ω = 0, green: Ω = 2*π*. (e) Simulated noiseless (top, red) x- and (bottom, blue) y-polarized images containing two emitters separated by distances of (left to right) 1 nm, 139 nm, 414 nm, and 620 nm. Magenta dot: center position for each emitter. (f-i) Detection rate, precision, and accuracy of Deep-SMOLM for estimating positions and orientations from images containing two emitters at various separations. (f) Deep-SMOLM (black) Jaccard index and the corresponding number of (orange solid) true-positive (TP), (orange dash) false-negative (FN), and (orange dot) false-positive (FP) emitters. (g) Deep-SMOLM (black) precision *σ*_*r*_ and (orange) accuracy *r* − *r*_0_ for estimating 2D position ***r***. (h) Deep-SMOLM (black) orientation precision *σ*_*ξ*_ and (orange) absolute mean orientation bias *ξ* (Eqn. 5). (i) Deep-SMOLM (black) precision *σ*_Ω_ and (orange) accuracy Ω − Ω_0_ for measuring wobble angle Ω.

We quantify Deep-SMOLM’s precision using simulated images of single emitters whose positions and orientations are random; for each orientation, 200 images are used to calculate the precision (Fig. 2(b-d) and Fig. S4). We use mean angular standard deviation *σ*_*δ*_ (MASD, Eqn. S7) to quantify the combined precision for measuring *θ* and *ϕ*. Deep-SMOLM gives an average 3D orientation estimation precision *σ*_*δ*_ of 3.8^◦^ and an average wobble angle precision *σ*_Ω_ of 0.32 sr for emitters with wobble angle Ω of 0 or 2 sr and with 1000 signal photons and 2 background photons detected per pixel. Deep-SMOLM also shows great performance for estimating the 2D position with an average precision of *σ*_*r*_ of 8.5 nm. For comparison, we compute the Cramér–Rao bound (CRB), which quantifies the best-possible estimation precision for any unbiased estimator [39]. For estimators with insignificant bias, CRB serves as a measure of optimal performance but may not be a true lower bound [40]. The precision for estimating the 3D orientation and 2D position is close to CRB precision (0% and 3% better than the CRB precision for 3D orientation and wobble angle, and 2% worse than the CRB precision for measuring 2D position) averaged over emitters with Ω = 0 sr and Ω = 2 sr, indicating the excellent optimum of Deep-SMOLM.

To quantify the bias *ξ* in mean orientation, we calculate the (non-negative) angular distance between the Deep-SMOLM’s estimated orientation 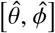 and the ground truth orientation [*θ, ϕ*], given by

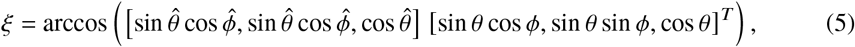

where the superscript *T* denotes a matrix-vector transpose. We note that Deep-SMOLM shows bias in estimating 3D orientations (an average mean orientation bias *ξ* of 1.3^◦^ and average wobble angle bias Ω − Ω_0_ of 0.13 sr, Fig. S4); this bias could enable Deep-SMOLM to achieve slightly better estimation precision than the CRB [40] (Fig. S4).

Robust single-molecule imaging, especially *in vivo*, necessitates an estimation algorithm that can reliably detect and estimate parameters from emitters whose images overlap [41, 42]. Early algorithms for measuring simultaneously SM orientations and positions either cannot cope with image overlap [9, 11, 23], are very computationally expensive [18], or can become stuck in local minima leading to correlated orientation and position biases [14]. To facilitate Deep-SMOLM’s robustness to these imaging conditions, we train it using simulated images containing both well-separated and overlapped DSFs corrupted by Poisson shot noise (SI section 4.ii).

We validate Deep-SMOLM’s ability to analyze overlapping images of SMs by simulating images containing 2 molecules separated by various distances (0-1000 nm) with fixed (Ω = 0) and random orientations and 1000 signal photons and 2 background photons detected per pixel. Deep-SMOLM is able to achieve a Jaccard index > 0.95 as long as the two emitters are separated by at least 139 nm, corresponding to an average 43% area overlap in their DSFs (Fig. 2(e,f), see SI section 3 for details on the Jaccard index and DSF overlap). More surprisingly, DeepSMOLM achieves accuracy and precision performance on par with non-overlapping emitters when analyzing images of emitters separated by just 414 nm, corresponding to 17% overlap in DSFs (Fig. 2(e-i), see Fig. S12 for performance at a lower signal-to-background ratio (SBR)).

### ii. 5D imaging of simulated biological fibers

To validate Deep-SMOLM for imaging a complex and densely labelled structure, we devised a synthetic structure containing nine 1D fibers as shown in Figs. 3(b) and S9. We designed the 3D orientations of the emitters to vary systematically with polar angles *θ* shown in Fig. 3(b) and azimuthal angles perpendicular to each fiber. In addition, all emitters are fixed in orientation (Ω = 0, see Fig. S14 for imaging emitters with wobble Ω = 2 sr). The emitters have a broad signal distribution with a mean of 1000 photons and 2 background photons detected per pixel (See Fig. S8(a) for photon distribution). Each simulated x- and y-polarized image (SI section 4) analyzed by Deep-SMOLM contains a random sampling of molecules with high activation density; each frame contains 7 to 15 emitters (the diameter of the structure is 2 *µ*m, see SI section 4.iii for details on image generation, and see Visualization 1 for simulated single molecule images). Importantly, the emitter density is lowest near the vertical midpoint of the structure, while a high density of emitters exists at the top and bottom “poles” where the fibers intersect.

**Fig. 3.**
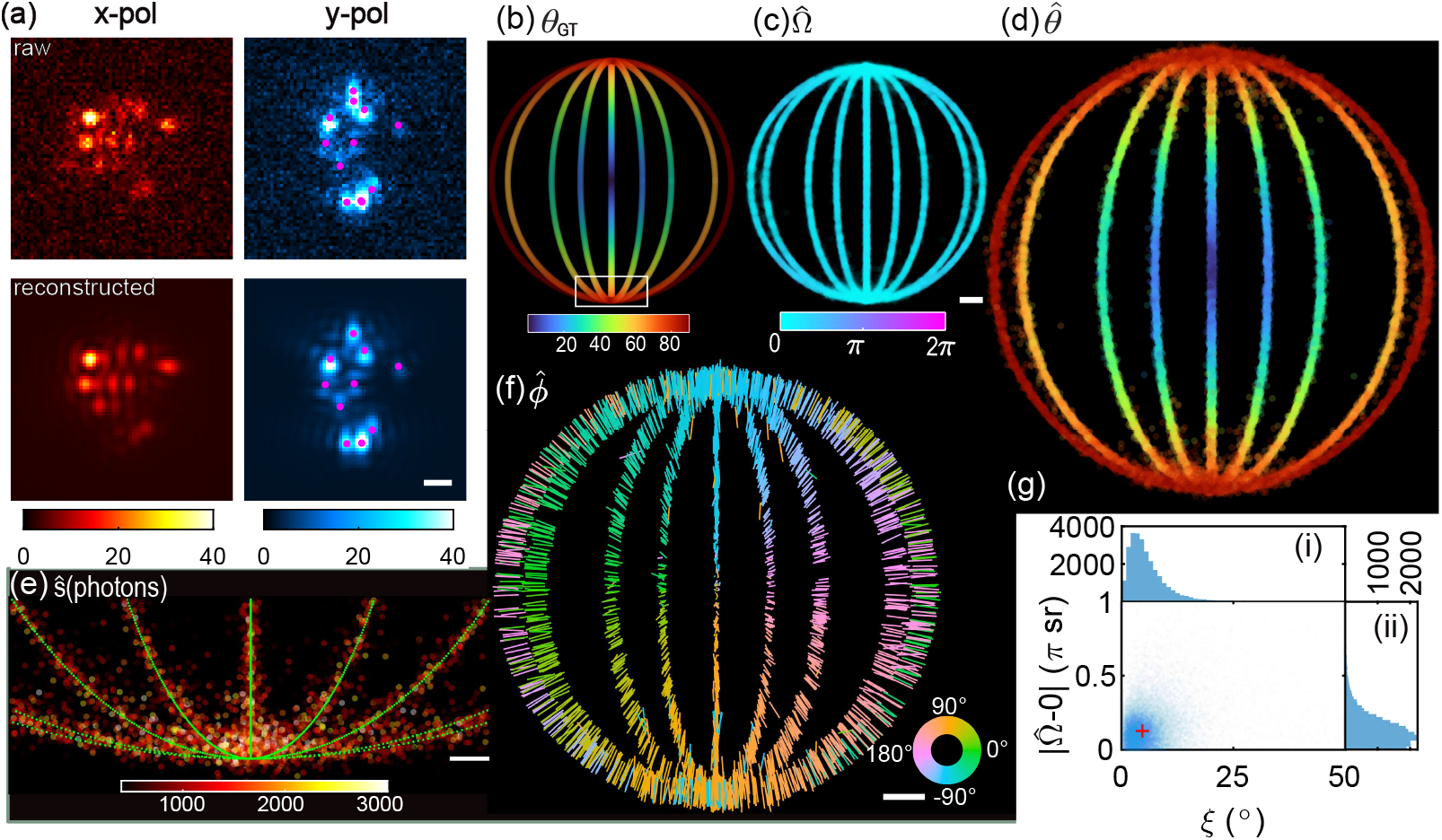
Deep-SMOLM 5D imaging using simulated data. (a)(top) Simulated raw image compared to (bottom) images reconstructed from Deep-SMOLM estimates. Magenta dots: center position of each SM. (b) Synthetic structure containing nine 1D fibers color-coded with the ground truth polar angle *θ*_GT_. (c) Deep-SMOLM measured wobble angle 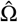 (ground truth Ω_GT_ = 0 sr). (d) Estimated polar angle 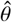. (e) Emitters within the white box shown in (b). Colormap: estimated signals *ŝ* (photons). (f) Estimated azimuthal angle 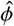, where the length and direction of each line depict the magnitude of the in-plane orientation sin 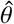 and direction of estimated azimuthal angle 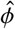, respectively. The ground truth orientations are perpendicular to the fibers. (g) Wobble angle estimation bias 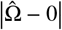 versus mean orientation estimation bias *ξ* (Eqn. 5). (Right) Distribution of wobble angle estimation bias and (top) mean orientation estimation bias. Scalebars: (c,f) 200 nm, (e) 50 nm.

Despite the extremely high degree of overlap in DSFs (Fig. 3(a)), Deep-SMOLM detects and localizes each emitter with excellent detection efficiency, achieving a 0.84 Jaccard index near the middle of the structure where fibers are more separated versus a 0.77 Jaccard index near the poles where DSF overlaps dominate (Fig. S13(a)). Deep-SMOLM readily resolves all nine fibers very well as shown in Fig. 3(e). However, Deep-SMOLM cannot resolve fibers <∼10 nm apart from each other due to the low SBR of the dataset; Deep-SMOLM achieves a mean spatial resolution of *σ*_*r*_ = 9.2 nm for an average of 1000 photons detected (Fig. S13(b)).

Deep-SMOLM’s 3D orientation estimates also match the ground truth very well (Fig. 3(c,d,f,g)). The median estimation bias in 3D orientation is *ξ* = 4.7^◦^, and the median bias in wobble is 0.13*π* sr, as expected from the impact of Poisson shot noise and measuring absolute (non-negative) bias in *ξ* and Ω [14, 43]. Furthermore, Deep-SMOLM achieves excellent orientation precision (a standard deviation *σ*_*ξ*_ in orientation bias of 7.1^◦^ and a wobble angle precision *σ*_Ω_ of 0.5 sr, Fig. 3(g) and Fig. S13(c,d)), despite frequent DSF overlaps in the raw data (Visualization 1).

### iii. Experimental 5D imaging of amyloid fibrils

Amyloid aggregates are linked to various pathological diseases, e.g., Parkinson’s and Alzheimer’s disease [44, 45]. Previous studies have shown that upon binding transiently to amyloid fibrils, Nile red (NR) orients itself parallel to their backbones [3,4,15]. Thus, imaging NR enables us to validate Deep-SMOLM’s performance for 5D SM imaging of orientations and positions from experimental data.

In typical biological imaging experiments, both optical aberrations and drift of the objective’s focal plane (FP) change affect the image of a SM. To mitigate microscope aberrations, we train Deep-SMOLM using DSFs calibrated [46] from experimental microscope images (SI section 5.i and Fig. S10). To mitigate the drift of FP in experimental data, we train Deep-SMOLM using simulated images of emitters captured with various FP positions *z* ∈ (−150, 150) nm, where *z* = 0 denotes focusing at the coverslip interface.

To validate Deep-SMOLM’s performance for analyzing images containing overlapped DSFs, we use NR at a high concentration, and thus an average high blinking rate of 0.6 emitters per µm^2^ (Visualization 2). Compared to diffraction-limited imaging (Fig. 4(b)), Deep-SMOLM’s reconstructed localization microscopy image resolves the network of intertwined amyloid fibrils with superior detail (Fig. 4(a)). Based on Fourier ring correlation [47], Deep-SMOLM attains a spatial resolution of *σ*_*r*_ = 11.0 nm (SI section 3.v and Fig. S6). Indeed, fibers spaced 55 nm apart are clearly resolved (Fig. 4(a) inset).

**Fig. 4.**
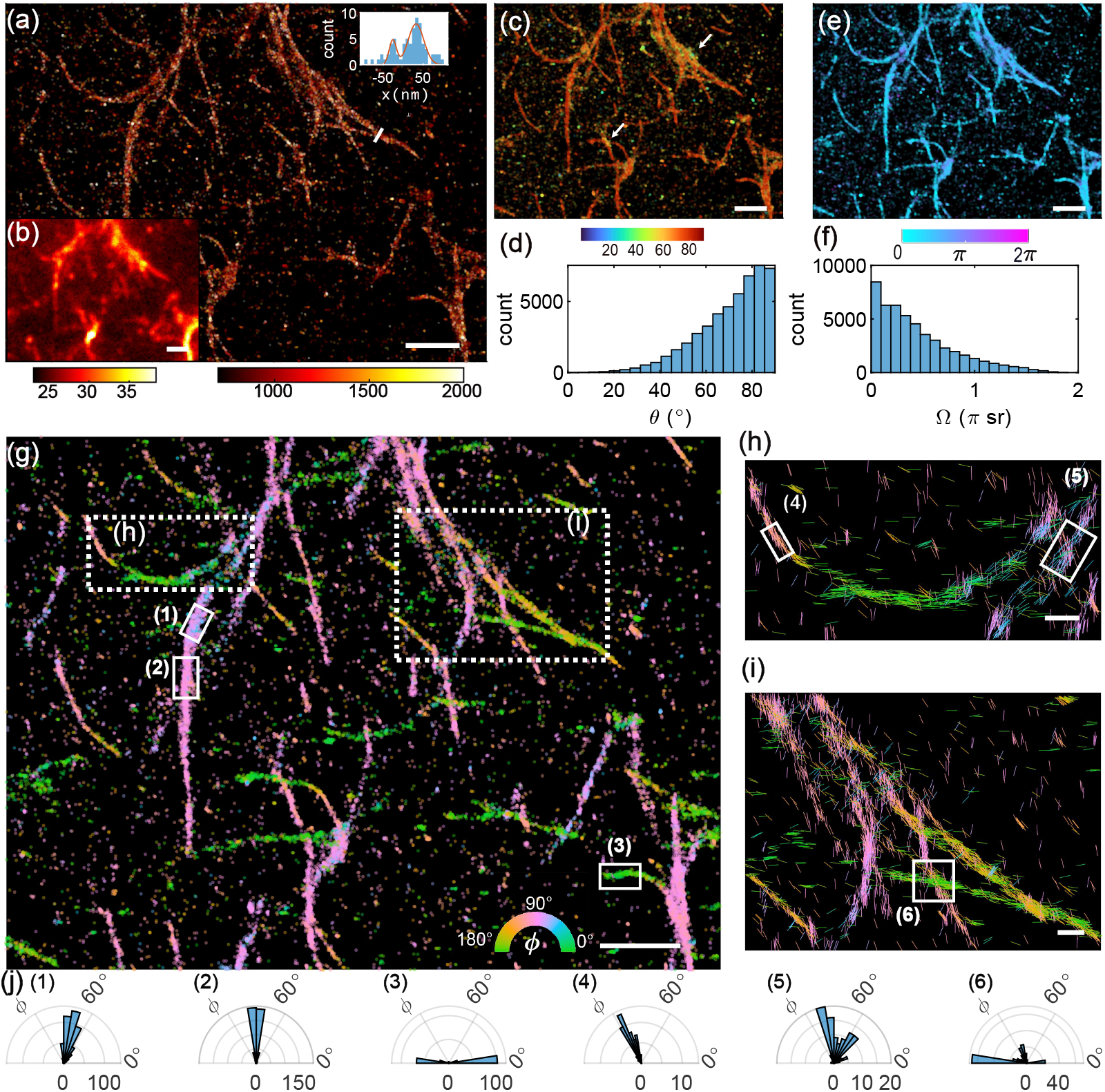
5D SMOLM images of Nile red (NR) transiently bound to A*β*42 amyloid fibrils. (a) SMLM reconstruction compared to (b) the corresponding diffraction-limited image. Colorbars: (a) signal photons for each detected emitter and (b) photons per 58.5 × 58.5 nm^2^ pixel. (a) Inset: distribution of molecule positions *x* along the white line in (a). Red line: double-Gaussian fit. (c) Spatial map (colorbar: deg) and (d) overall distribution of NR polar angles 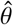. (e) Spatial map (colorbar: sr) and (f) overall distribution of NR wobble angles 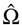. (g) Spatial map of azimuthal angles 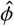 for NR with polar angle 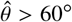 and wobble angle Ω < 2 sr. Each SM is represented as a 1 nm filled circle in (a,c,d) and represented as a 2 nm filled circle in (g). (h,i) All NR positions and orientations detected within the dotted white boxes in (g), depicted as line segments. Their lengths and directions indicate the magnitude of their in-plane orientations (sin(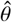)) and their azimuthal orientations 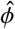, respectively. Colorbar: 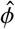 (deg). (j) Azimuthal angle 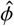 distribution of all NR molecules with each solid white box in (g-i). Scale bars: (a-c,e,g) 1 µm, (h,i) 200 nm.

Deep-SMOLM’s orientation estimates show that 75% of NR’s polar orientations 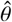 are greater than 60^◦^, i.e., parallel to the covership (Fig. 4(c,d)). However, when bound to dense entangled fibrils, some NR molecules have smaller polar angles 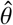 as indicated by arrows in Fig. 4(c), which indicates that fibrils are tilted out of the coverslip plane when they overlap with one another. In addition, 72% of NR molecules have a wobble angle Ω smaller than 2 sr, indicating that they rigidly bind to fibrils (Fig. 4(e,f)).

Examining Deep-SMOLM’s measurements of NR azimuthal angles 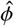, we find that NR is well-aligned with the long axis of each amyloid fibril (Fig. 4(g-j)); the average orientation of NR varies smoothly as the amyloid fibrils bend and curve as shown in Figs. 4(g-j), especially in Fig. 4(h). Within crowded regions containing entangled fibrils, Deep-SMOLM accurately resolves NR orientations aligned with each individual fiber, as shown in Fig. 4(h,i); accordingly, multiple peaks are present in the histograms of azimuthal angles 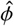 (Fig. 4(j)(5-6)). Notably, DSFs located at these entangled regions are highly overlapped but still accurately estimated (Visualization 3).

## 4. Discussion and conclusion

Here, we demonstrate a deep learning-based estimator, called Deep-SMOLM, for simultaneously estimating 3D orientations and 2D positions of single molecules from a microscope implementing an engineered dipole-spread function [14]. Compared to traditional optimization approaches, Deep-SMOLM achieves superior estimation precision for both 3D orientation and 2D position that is on average within 3% of the best-possible precision (Fig. 2(b-d)). In general, designing a loss function for high-dimensional estimation and balancing weights among multiple parameters in particular, are always challenging. We attribute the superior performance of Deep-SMOLM to the linearity of estimating brightness-weighted orientational second moments from noisy SM images (Fig. 1(c) and Eqn. 2); otherwise, directly estimating orientation angles [*θ, ϕ*, Ω] is ill-conditioned and unstable. Importantly, for high-performance DSFs, each brightness-weighted orientational moment contributes approximately equally to the final DSF shape [12, 14]. Further, we have designed 3D orientations and 2D positions to be orthogonally encoded into the intensities and spatial positions, respectively, of Gaussian spots within Deep-SMOLM’s output images (Fig. 1(c)). These design strategies make the tuning of weights among the six output images unnecessary for training Deep-SMOLM (SI section 2.ii), and the resulting training among 5-dimensional estimates is well-balanced (Fig. S2).

As demonstrated on both simulated structures (Fig. 3) and experimental amyloid fibrils (Fig. 4), Deep-SMOLM shows excellent performance for estimating overlapped DSFs (Fig. 2(e-i)), e.g., at a density of ∼0.6 emitter/µm^2^, which is ∼2 times denser than that allowed by traditional optimization-based algorithms. This capability should allow Deep-SMOLM to achieve a ∼2× speed-up in SMOLM data acquisition by enabling fluorescent probes to blink at higher rates and to be used at higher concentrations. Moreover, Deep-SMOLM requires relatively little training time and data (∼2 h and 30,000 noisy images containing ∼330,000 total emitters). Once trained, Deep-SMOLM estimates 3D orientations and 2D positions ∼10 times faster on a per-frame basis than iterative algorithms like RoSE-O [18] (∼30 seconds for Deep-SMOLM, ∼6 minutes for RoSE-O for analyzing 1,000 frames). Taking into account the fewer raw frames needed to compute reliable SMOLM reconstructions, Deep-SMOLM exhibits an overall 20× speed-up in computation time to obtain the same total number of localizations versus iterative algorithms. We note that correlations in SM blinking between frames [48] can also be used to further enhance Deep-SMOLM’s performance for estimating overlapped emitters. We anticipate that Deep-SMOLM will enable fast high-dimensional SMOLM imaging of dynamic processes and potentially discover structural changes on the sub-minute timescale.

In the near future, we plan to extend Deep-SMOLM to simultaneously estimate the 3D positions and 3D orientations of overlapping molecules; this analysis is inherently much more complex than 2D SMOLM due to the strong sensitivity of pixOL’s basis images to the axial position of each emitter [14]. Further, especially when imaging *in vivo* cellular structures in 3D, estimators need improved robustness against model mismatch, e.g., accommodating non-uniform background and field- and depth-dependent optical aberrations [13, 49]. However, exhaustively training a network to anticipate all practical perturbations of an imaging system is extremely challenging. Thus, simultaneously learning model mismatch together with estimating 3D position and 3D orientation could enhance Deep-SMOLM’s performance in challenging imaging conditions. This adaptive approach may be key to ensuring that networks are sufficiently generalizable for *in vivo* super-resolution imaging and could unlock the full potential of SMOLM for cellular and tissue-scale imaging.

## Supporting information

visualization 1

visualization 2

visualization 3

## Funding

National Science Foundation (NSF) (ECCS-1653777); National Institute of General Medical Sciences (R35GM124858).

## Acknowledgments

The authors thank Hesam Mazidi, Joseph O’Sullivan, Joseph Culver, Carlos Fernandez-Granda, and Qing Qu for helpful suggestions and comments. We are also grateful to Tianben Ding for help with amyloid aggregation. Amyloid-*β* peptides were synthesized and purified by Dr. James I. Elliott (ERI Amyloid Laboratory, Oxford, CT).

## Disclosures

The authors declare no conflicts of interest.

## Data availability

The Deep-SMOLM algorithm, forward model, training data, validation data, and experimental data are available via OSF [50], Github [51], and by request.

## Supplemental document

See Supplement 1 for supporting content.

## Notes

https://github.com/Lew-Lab/Deep-SMOLM

### Competing Interest Statement

The authors have declared no competing interest.

