## supplemental information for "Deep-SMOLM: Deep Learning Resolves the 3D Orientations and 2D Positions of Overlapping Single Molecules with Optimal Nanoscale Resolution"

This document provides supplementary information to "Deep-SMOLM: Deep Learning Resolves the 3D Orientations and 2D Positions of Overlapping Single Molecules with Optimal Nanoscale Resolution," offering details on the forward model, estimator architecture, performance quantification, and sample preparation.

### 1. RELATIONSHIP BETWEEN ORIENTATIONAL SECOND MOMENTS AND ORIENTATION ANGLES

The orientation of a dipole-like emitter can be represented by a unit vector $[\mu_x, \mu_y, \mu_z]$ or equivalent polar and azimuthal angles $[\theta, \phi]$ in spherical coordinates, where $[\mu_x, \mu_y, \mu_z] = [\sin(\theta)\cos(\phi), \sin(\theta)\sin(\phi), \cos(\theta)]$. If an emitter "wobbles" through a range of directions represented by a hard-edged cone with solid angle $\Omega$ within a camera's exposure time, then its orientational trajectory can be represented using orientational second moments $\boldsymbol{m} = [\langle\mu_x^2\rangle, \langle\mu_y^2\rangle, \langle\mu_z^2\rangle, \langle\mu_x\mu_y\rangle, \langle\mu_x\mu_z\rangle, \langle\mu_y\mu_z\rangle]^T \in \mathbb{R}^6$ as given by

$$\langle\mu_x^2\rangle = \gamma\mu_x^2 + (1-\gamma)/3, \tag{S1a}$$

$$\langle\mu_y^2\rangle = \gamma\mu_y^2 + (1-\gamma)/3, \tag{S1b}$$

$$\langle\mu_z^2\rangle = \gamma\mu_z^2 + (1-\gamma)/3, \tag{S1c}$$

$$\langle\mu_x\mu_y\rangle = \gamma\mu_x\mu_y, \tag{S1d}$$

$$\langle\mu_x\mu_z\rangle = \gamma\mu_x\mu_z, \tag{S1e}$$

$$\langle\mu_y\mu_z\rangle = \gamma\mu_y\mu_z, \text{ and} \tag{S1f}$$

$$\gamma = 1 - \frac{3\Omega}{4\pi} + \frac{\Omega^2}{8\pi^2}, \tag{S1g}$$

where $\langle\cdot\rangle$ denotes a temporal average over the camera acquisition period. The rotational constraint $\gamma$ and the solid angle $\Omega$ are equivalent ways to quantify an emitter's rotational diffusion [1].

### 2. DEEP-SMOLM PROCESSING ARCHITECTURE

#### i. Network structure

We build and optimize our neural network using Pytorch. Our neural network structure (Fig. S1) is adapted from DeepSMOLM3D [2]. More details are located in Fig. S1 and within the shared code in [3, 4].

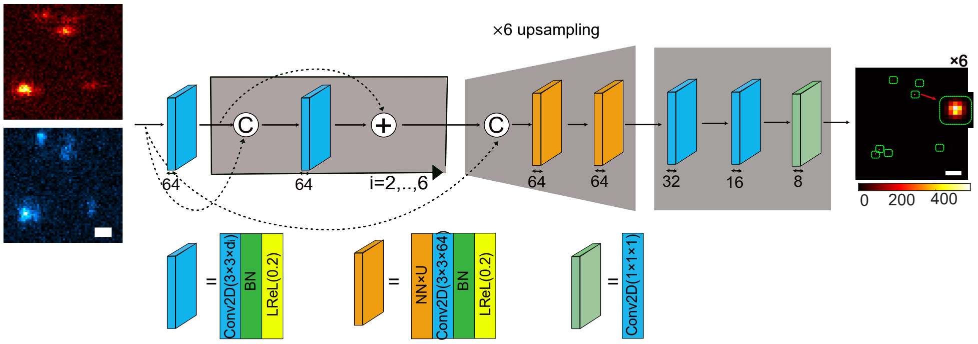

**Fig. S1.** Neural network architecture. A set of (top, red) x- and (bottom, blue) y-polarized images $\boldsymbol{I}_{\text{in}}$ is first passed through a batch normalization (BN) to normalize pixel values. Next, the normalized image $\boldsymbol{I}_{\text{norm}}$ is passed to the fully convolutional architecture composed of three parts. In the first section, the normalized image $\boldsymbol{I}_{\text{norm}}$ goes through six (blue box) dilated convolution blocks. The dilated convolution blocks have a fixed number of channels (64) comprising a 2D convolutional layer (Conv2D), a batch normalization (BN) layer, and a LeakyRELU (LReL) activation function with a negative slope of 0.2. The 2D convolutional layer has a filter size of $3 \times 3$ and dilution rate of $d_i$, where the index $i$ denotes the $i^{\text{th}}$ dilated convolution block, $d_1 = d_2 = d_5 = d_6 = 1$, $d_3 = 2$, and $d_4 = 4$. Concatenation (C) and element-wise addition (+) are used to improve the gradient flow. The second part of the network is designed to upsample the image laterally by a factor of 6 by using one $3\times$ (orange box) resize-convolution block and one $2\times$ resize-convolution block. NN×U: U× upsampling operator. The third part of the network creates output images by gradually reducing the number of channels to a final output of six, corresponding to the six output images (Fig. S3). The (blue box) dilated convolution blocks in the third part of the network have a filter size of $3 \times 3$ and dilution rate of 1.

#### ii. Training the neural network

Considering the memory of our GPU, we use a batch size of 32 images for training the neural network. We use stochastic gradient descent (SGD) to optimize the neural network. The learning rate is set to be 0.001 at the beginning. If the validation loss doesn't decrease for three consecutive epochs, then the learning rate is reduced by a factor of 0.1. This strategy guarantees an optimal learning rate even if the learning rate is too large at the beginning. We use momentum factor of 0.9, and a weight decay of 0.0005 for SGD.

Since we designed 3D orientations and 2D positions to be orthogonally encoded into the intensities and spatial positions, respectively, of Gaussian spots within Deep-SMOLM's output images (Fig. 1(c)), there is no need to balance the contributions of 3D orientation estimation errors versus 2D position estimation errors in the loss function. The intensity distribution of the DSF is linearly proportional to the orientational moments as shown in Eqn. 1. Each brightness-weighted orientational moment contributes approximately equally to the final DSF shape (Fig. S7). Without tuning the weights among six images (Eqn. 4), the training loss from the six brightness-weighted orientational moment are already at the same scale, indicating well-balanced weights among the six output images (Fig. S2(a)).

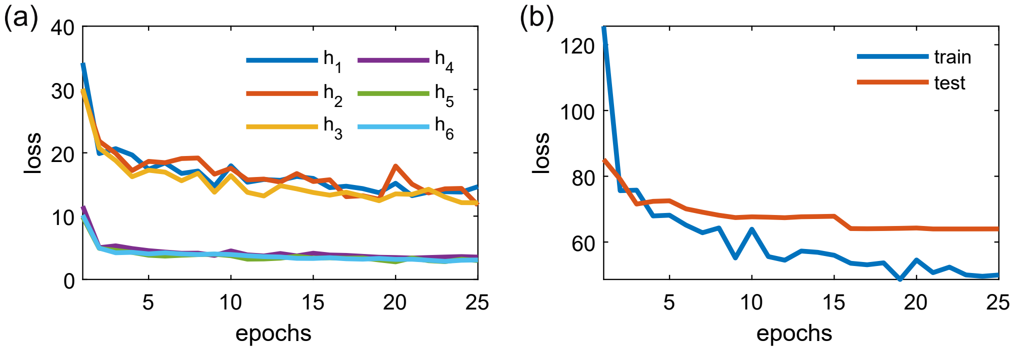

**Fig. S2.** Training losses of Deep-SMOLM. (a) Training losses of Deep-SMOLM for each brightness-weighted orientational moment image $\boldsymbol{h}_l$. (b) Training loss (Eqn. 4) and testing loss versus training epoch.

##### iii. Postprocessing algorithm

We design a postprocessing algorithm to compile the six output images from the network (Fig. S3) into a list of SMs, each with a measured 2D position $\hat{\boldsymbol{r}}$, intensity $\hat{s}$, and 3D orientation $[\hat{\theta}, \hat{\phi}, \hat{\Omega}]$. Since each detected emitter is represented using a 2D Gaussian spot co-located within each of the 6 output images $\boldsymbol{h}_r$, the postprocessing algorithm uses a pattern-matching algorithm to find Gaussian patterns within the sum of the first three images output from the network, given by $\hat{\boldsymbol{h}}_1(\boldsymbol{r}) + \hat{\boldsymbol{h}}_2(\boldsymbol{r}) + \hat{\boldsymbol{h}}_3(\boldsymbol{r})$. This summed image represents the brightnesses (signal photons) and positions of all molecules detected by the network.

For each detected Gaussian pattern, one $7 \times 7$ images $\bar{\boldsymbol{h}}_l$ is cropped from each of the six images $\hat{\boldsymbol{h}}_l(\boldsymbol{r})$ output by the network, where each is centered at the brightest pixel $[x_0, y_0]$. A threshold of 200 photons is used to filter out emitters with low signal photons. The signal of each emitter is calculated as

$$\hat{s} = \frac{1}{A} \sum_{p=1}^{7} \sum_{k=1}^{7} \sum_{l=1}^{3} \bar{\boldsymbol{h}}_l^{k,p}, \tag{S2}$$

where $[\cdot]_l^{k,p}$ represents the pixel on the $k^{\text{th}}$ row and the $p^{\text{th}}$ column of the $l^{\text{th}}$ cropped image $\bar{\boldsymbol{h}}$ and $A$ is the summed intensity of Gaussian kernel used in creating the ground truth image $\boldsymbol{h}_l(\boldsymbol{r})$.

A simple centroid estimator is used to calulate the position of each emitter as

$$\hat{x} = E\left[x_0 + \frac{1}{A\hat{s}} \sum_{p=1}^{7} \sum_{k=1}^{7} (p-4) \sum_{l=1}^{3} \bar{\boldsymbol{h}}_l^{k,p}\right] \text{ and} \tag{S3}$$

$$\hat{y} = E\left[y_0 + \frac{1}{A\hat{s}} \sum_{p=1}^{7} \sum_{k=1}^{7} (k-4) \sum_{l=1}^{3} \bar{\boldsymbol{h}}_l^{k,p}\right], \tag{S4}$$

where $E$ is the pixel size of output images (9.75 nm) and $[x_0, y_0]$ is pixel index of the center of the cropped image with respect to the original image. The orientational moments are calculated from the brightness of six Gaussian patterns as

$$m_l = \frac{1}{A\hat{s}} \sum_{p=1}^{7} \sum_{k=1}^{7} \bar{\boldsymbol{h}}_l^{k,p}. \tag{S5}$$

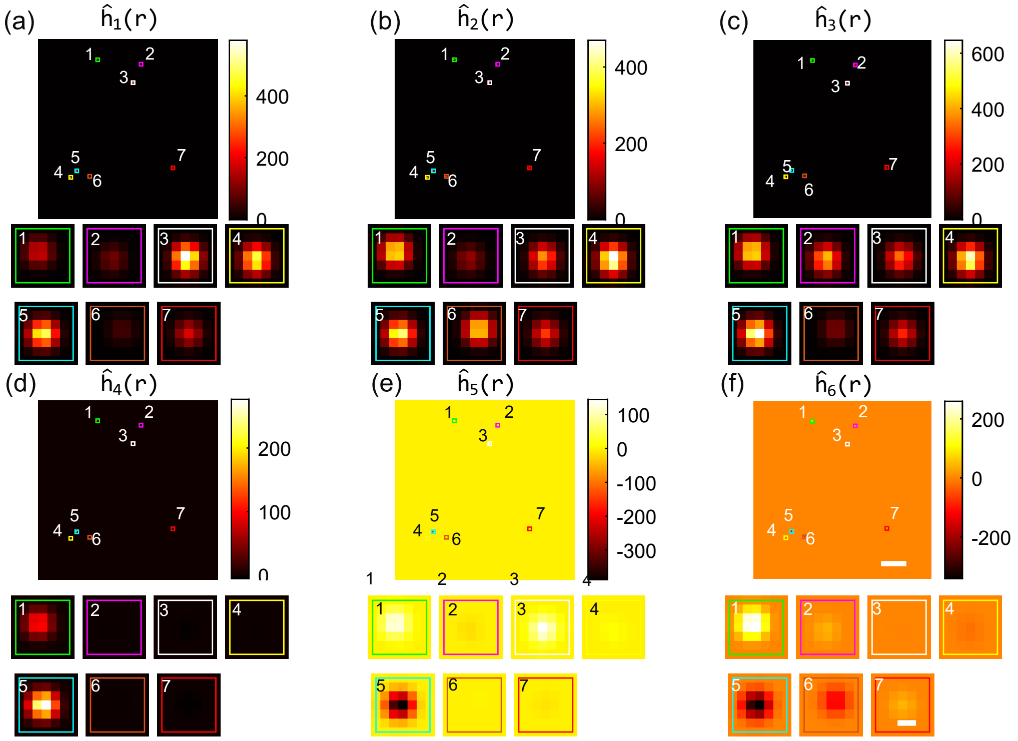

**Fig. S3.** (a-f) Six Gaussian-blurred brightness-weighted orientational second-moment images $\hat{\boldsymbol{h}}_l(\boldsymbol{r})$ output by Deep-SMOLM. Colorbars: intensity (a.u.). (1-7) Six $7 \times 7$ images $\bar{\boldsymbol{h}}_l$ cropped from $\hat{\boldsymbol{h}}_l(\boldsymbol{r})$, each corresponding to one of the 7 detected emitters. Scale bars: (a-f) 500 nm, (1-7) 20 nm.

The estimated second-moment vectors $\hat{\boldsymbol{m}}_q$ are next projected to first-moment orientation space $[\hat{\phi}, \hat{\theta}, \hat{\Omega}]$ using a weighted least-square estimator as follows:

$$[\hat{\phi}, \hat{\theta}, \hat{\Omega}] = \underset{\{\phi', \theta', \Omega'\}}{\arg\min} \left(\boldsymbol{m} - \boldsymbol{m}(\phi', \theta', \Omega')\right)^T \mathbf{F} \left(\boldsymbol{m} - \boldsymbol{m}(\phi', \theta', \Omega')\right). \tag{S6}$$

Note that weighting by the Fisher information matrix $\mathbf{F}$ ensures that more weight is given to the second moments $m_l$ for which pixOL demonstrates superior precision [5].

#### 3. QUANTIFYING THE PERFORMANCE OF DEEP-SMOLM

##### i. Mean angular standard deviation

To quantify the precision of estimating the mean orientation $[\theta, \phi]$, we calculate the mean angular standard deviation $\sigma_\delta$, which is the half-angle of the uncertainty cone for estimating the mean orientation direction [6]. It is a summary metric that combines the precision $\sigma_\theta$ of measuring $\theta$ and the standard deviation $\sigma_\phi$ of measuring $\phi$, given by

$$\sigma_\delta = 2 \arcsin \left( \sqrt{\frac{\sin(\theta)\sigma_\theta \sigma_\phi}{\pi}} \right), \tag{S7}$$

##### ii. Jaccard index

To compare estimates to the ground truth, we have to pair each estimated emitter with a ground-truth emitter. We choose to pair these emitters by minimizing the overall 2D Euclidean distance between matched points, as given by

$$[\hat{x^r}, \hat{y^r}] = \underset{\{\hat{x^r}, \hat{y^r}\}}{\arg\min} \sum_i \left[ (\hat{x}_i^r - x_i)^2 + (\hat{y}_i^r - y_i)^2 \right], \tag{S8}$$

where $[\hat{x}_i^r, \hat{y}_i^r]$ is the 2D position of the $i^{\text{th}}$ estimated emitter reordered based on the order of matched ground-truth emitters, $[x_i, y_i]$ is the ground truth position of the $i^{\text{th}}$ estimated emitter. After the matching, estimated emitters outside a threshold distance of 150 nm from the ground truth are designated as false positives (FP).

The Jaccard index is used to quantify the detection accuracy of an estimator. We calculate the Jaccard index as

$$\text{Jaccard} = \frac{\text{TP}}{\text{FN} + \text{FP} + \text{FP}}, \tag{S9}$$

where true positive (TP) is defined as a detected emitter that has a corresponding ground truth emitter, false negative (FN) is defined as a missing emitter, and false positive (FP) is defined as a detected emitter that doesn't have a corresponding ground truth emitter.

##### iii. Overlapping percentage of two DSFs

To calculate the overlapping percentage of two overlapping DSFs, we first convert the DSF $\boldsymbol{I} \in \mathbb{R}^{56\times112}$ of each emitter to a binary image $\boldsymbol{I}^b$ as

$$\boldsymbol{I}^b = \begin{cases} 1, & \text{if } I_i \geq 0.05 \max(\boldsymbol{I}) \\ 0, & \text{if } I_i < 0.05 \max(\boldsymbol{I}) \end{cases}, \tag{S10}$$

where $[\cdot]_i$ represents the $i^{\text{th}}$ pixel of the $\boldsymbol{I}$, $\max(\cdot)$ is the maximum operator. The overlapping percentage $O$ is calculated based on the overlapping of binary image $\boldsymbol{I}^b$ of two emitters using

$$O = \frac{2\sum_i I^b_{1,i} I^b_{2,i}}{\sum_i I^b_{1,i} + \sum_i I^b_{2,i}} \times 100\%, \tag{S11}$$

where $[\cdot]^b_{q,i}$ represents the $i^{\text{th}}$ pixel of the binary image $\boldsymbol{I}^b_q$ for the $q^{\text{th}}$ emitter.

##### iv. Estimation bias of Deep-SMOLM

We note that Deep-SMOLM estimates exhibit a small but non-negligible bias (Fig. S4). In terms of the estimated orientational second moments $\boldsymbol{m}$, the bias skews towards the median value of $\boldsymbol{m}$ (Fig. S5). To improve accuracy, one can correct the bias of the orientational second moments in postprocessing. However, we did not implement any bias corrections due to the relatively small bias magnitude compared to estimation precision, caused by relatively dim emitters in our SMOLM measurements.

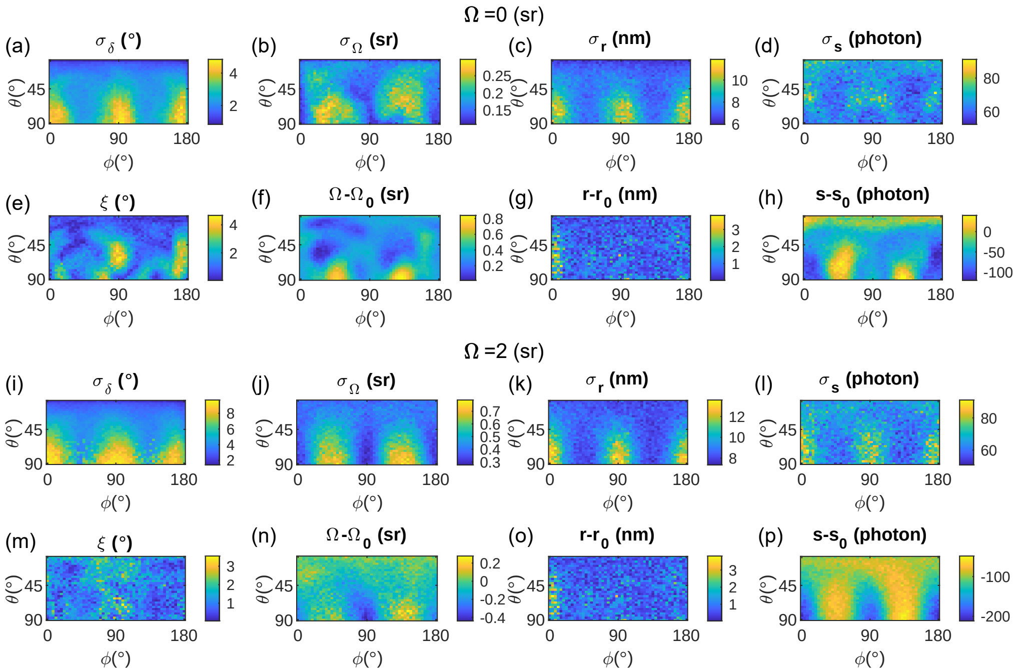

**Fig. S4.** Deep-SMOLM estimation precision and accuracy. (a-d,i-l) Precision and (e-h,m-p) accuracy for emitters with wobble angle $\Omega$ of (a-h) 0 and (i-p) 2 sr. (a,i) Mean angular standard deviation $\sigma_\delta$, (b,j) wobble angle precision $\sigma_\Omega$, (c,k) position estimation precision $\sigma_r$, and (d,l) intensity estimation precision $\sigma_s$. (e,m) (Non-negative) angular distance $\xi$ between the estimated orientation $[\hat{\theta}, \hat{\phi}]$ and ground truth orientation $[\theta, \phi]$ (Eqn. 5), (f,n) wobble angle bias $\Omega - \Omega_0$, (g,o) 2D position bias $r - r_0$, and (h,p) signal photon estimation bias $s - s_0$. At each orientation, 200 independent images were generated for emitters with 1000 signal photons in total and 2 background photons per pixel detected. The 3D orientations and 2D positions are estimated using Deep-SMOLM.

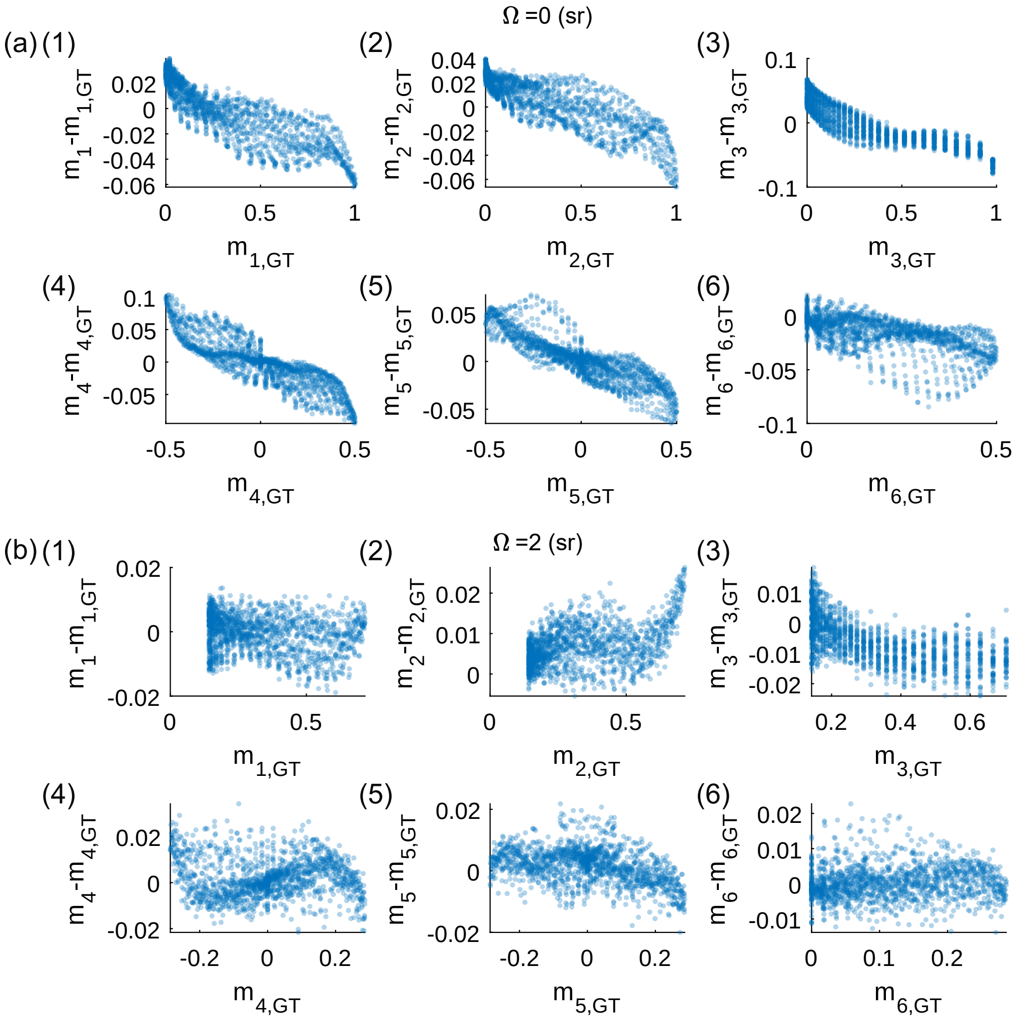

**Fig. S5.** Deep-SMOLM estimation bias of the orientational second moments $\boldsymbol{m}$ for emitters with wobble angle $\Omega$ of (a) 0 and (b) 2 sr. (1-6) Mean orientational second moment estimation bias $m_l - m_{l,\mathrm{GT}}$ versus the ground truth $m_{l,\mathrm{GT}}$ for the $l^{\mathrm{th}}$ orientational moment $m_l$. Each scatter point is an orientation shown in Fig. S4. The bias is averaged over 200 independent images.

##### v. Fourier ring correlation

We calculate the spatial resolution $\sigma_r$ of experimental SMOLM images using Fourier ring correlation (FRC) as described in [7]; see Fig. S6 for the spatial resolution of the amyloid fibril SMOLM data in Fig. 4. Emitters with signal photons larger than 600 are randomly divided into two subsets. We generate standard SMLM images $f_1(r)$ and $f_2(r)$ from the two subsets by binning localizations using a pixel size of 5 nm; the value of each pixel represents the number of emitters located within that pixel. We then compute the Fourier transforms $\hat{f}_1(v)$ and $\hat{f}_2(v)$ of the two images and discretize Fourier space into multiple rings. Each ring $\mathcal{C}$ represents a spatial frequency. FRC is calculated using the correlation of $\hat{f}_1(v)$ and $f_2\hat{(v)}$ over pixels within the ring $\mathcal{C}$ as

$$\mathrm{FRC}(\mathcal{C}) = \frac{\sum_{v\in\mathcal{C}} \hat{f}_1(v)\hat{f}_2^{\,*}(v)}{\sqrt{\sum_{v\in\mathcal{C}} \left\|\hat{f}_1(v)\right\|^2 \sum_{v\in\mathcal{C}} \left\|\hat{f}_2(v)\right\|^2}}, \tag{S12}$$

where $(\cdot)^*$ represents the complex conjugate operator. To determine resolution, we use the $2\sigma$ curve as a threshold, given by

$$F_{2\sigma}(\mathcal{C}) = \sqrt{\frac{8}{N_p(\mathcal{C})}}, \tag{S13}$$

where $N_p(C)$ represent the number of pixels within ring $\mathcal{C}$. We then find the spatial frequency $v_c$ of the first crossing between $\mathrm{FRC}(\mathcal{C})$ and $F_{2\sigma}(\mathcal{C})$. The resolution $\sigma_r$ is calculated as

$$\sigma_r = \frac{1}{2v_c}. \tag{S14}$$

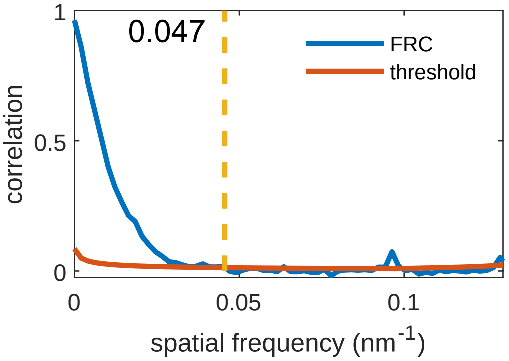

**Fig. S6.** Fourier ring correlation (FRC) of the amyloid fibril SMOLM data shown in Fig. 4. Blue: FRC $\mathrm{FRC}(\mathcal{C})$; red: threshold $F_{2\sigma}(\mathcal{C})$; yellow dash: first intersection between $\mathrm{FRC}(\mathcal{C})$ and $F_{2\sigma}(\mathcal{C})$.

#### 4. FORWARD MODEL FOR GENERATING SYNTHETIC DATA

##### i. Forward model

The image of an SM produced by the microscope equals the sum of basis images $\mathbf{B}$ weighted by the orientational second moments as shown in Eqn. 1. The $l^{\text{th}}$ basis matrix $\mathbf{B}_l$ is a concatenated image of x- and y-polarized basis images, corresponding to the system's response to the $l^{\text{th}}$ orientational second moment (Fig. S7). To accurately generate images containing $Q$ emitters at various positions $\boldsymbol{r}_q$ (Eqn. 2), the basis images $\boldsymbol{B}(\boldsymbol{r}-\boldsymbol{r}_q)$ centered at arbitrary continuous 2D positions $\boldsymbol{r}_q$ must be computed. To reduce the computational burden, we use the *imtranslate* function in MATLAB to calculate $\boldsymbol{B}(\boldsymbol{r}-\boldsymbol{r}_q)$ via bi-cubic interpolation.

After generating the noiseless images using Eqn. 2, Poisson shot noise is added to each pixel independently as

$$\boldsymbol{I}^p = \mathrm{Poiss}(\boldsymbol{I}), \tag{S15}$$

where $\mathrm{Poiss}(\lambda)$ corresponds to generating random values from a Poisson distribution with mean of $\lambda$.

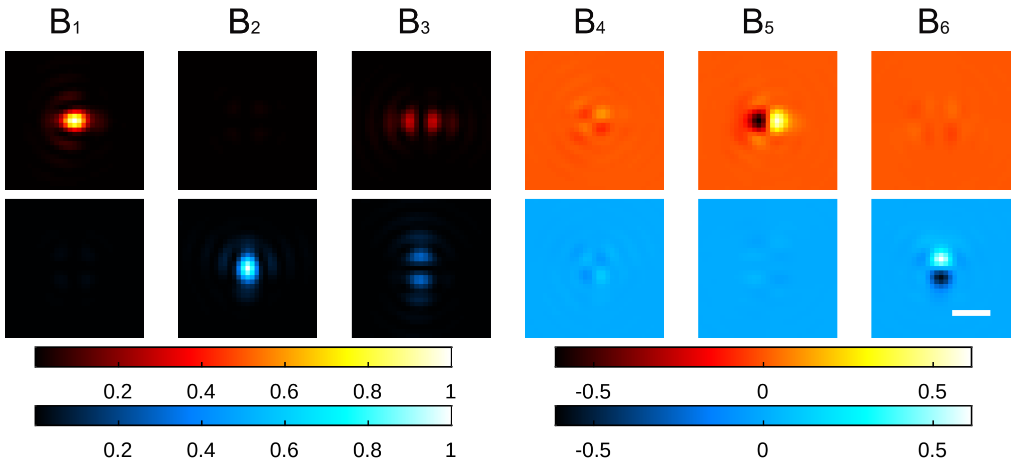

**Fig. S7.** Image plane basis images $\mathbf{B}_l$ corresponding to a polarization-sensitive imaging system (red: x-polarized, blue: y-polarized) with a pixOL phase mask [5] for an in-focus emitter. The image intensities are normalized relative to the brightest basis image ($\mathbf{B}_1$). Colorbars: normalized intensity. Scalebar: 500 nm.

#### ii. Generating training data

For one training image, the number of emitters $Q$ is drawn from a uniform distribution of $Q \in \{7, 8, .., 15\}$. A 2D position is randomly assigned to each emitter. The mean orientation $\boldsymbol{\mu} = [\mu_x, \mu_y, \mu_z]^T$ of each emitter is randomly generated using [8]

$$\mu_x = 2x_1\sqrt{1 - x_1^2 - x_2^2} \tag{S16a}$$

$$\mu_y = 2x_2\sqrt{1 - x_1^2 - x_2^2} \tag{S16b}$$

$$\mu_z = 1 - 2(x_1^2 + x_2^2), \tag{S16c}$$

where $x_1$, $x_2$ are uniformly distributed within $(-1, 1)$ and points for which $x_1^2 + x_2^2 \geq 1$ are rejected. The wobble angle $\Omega$ are computed by drawing $\gamma$ from a linear distribution (Eqn. S1). The signal photons of emitters have a broad distribution with mean of 1000 as show in Fig. S8(a), and all training images have mean background of 2 photons per pixel.

We generate 30K noiseless images based on the forward model (SI section 4.i) with signal and orientation parameters distributed as shown in Fig. S8. At each training epoch, we add Poisson shot noise to the noiseless images (Eqn. S15) and then pass the noisy images to Deep-SMOLM. We use 90% percent of the data for training, and remaining 10% as testing data.

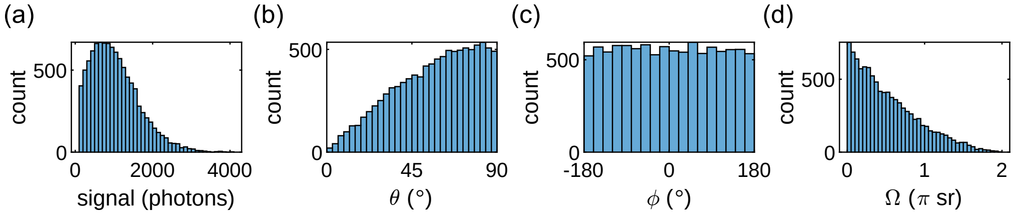

**Fig. S8.** (a) Signal $s$, (b) polar angle $\theta$, (c) azimuthal angle $\phi$, and (d) wobble angle $\Omega$ distributions for the 30K images used to train Deep-SMOLM. The signal distribution in (a) is also used for generating the simulated biological fibers in Fig. 3.

#### iii. Generating simulated biological fibers

The designed structure is used to initialize the 2D position of each emitter, and the 3D orientation of each is computed based on its position within the structure (Fig. S9). The signal of each emitter and background of each simulated image have distributions similar to the training data (Sec. 4.ii and Fig. S8(a)).

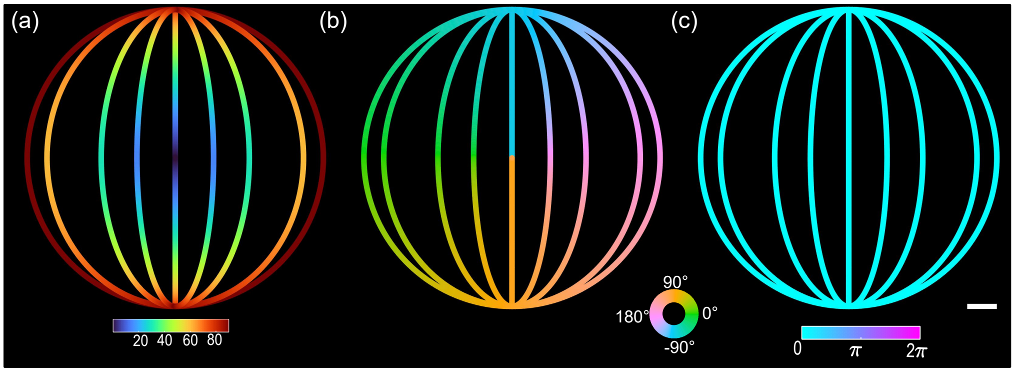

**Fig. S9.** Model structure of 1D fibers. (a) Ground truth polar angle $\theta$, (b) azimuthal angle $\phi$, and (c) wobble angle $\Omega$ for each emitter along the structure.

#### 5. EXPERIMENTAL IMAGING OF AMYLOID FIBRILS

##### i. Microscope calibration

We use the perfect pixOL phase mask (Fig. S10(a)) for generating all synthetic data used in Fig. 2 and Fig. 3. However, to improve pixOL DSF performance in our microscope [5], we use the conjugate pixOL phase mask (pixOL*, Fig. S10(b)) to collect and analyze experimental amyloid fibril data (Fig. 4).

We calibrate the imaging model to the imaging system's DSF by using fluorescent beads (100-nm diameter red 580/605 FluoSpheres, Invitrogen F8801). A phase-retrieval algorithm [9] is used to retrieve the experimental phase mask. To accurately characterize optical aberrations, the phase masks of the two polarized channels are estimated independently (Fig. S10(c,d)). These are consequently used to simulate images collected by each polarized channel for training Deep-SMOLM (Sec. 4.ii).

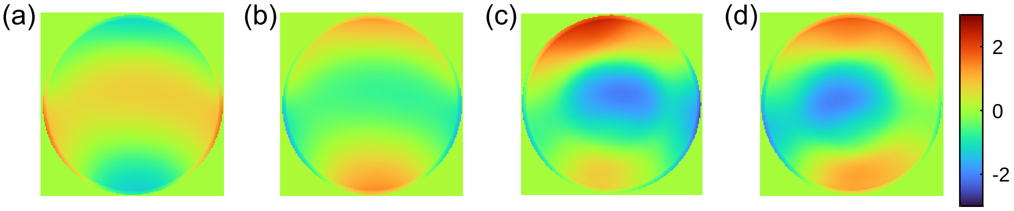

**Fig. S10.** Calibration of the conjugate pixOL phase mask pixOL* [5]. (a) Perfect pixOL phase mask. (b) Perfect conjugate pixOL* of the pixOL phase mask. (c,d) Calibrated experimental pupil phase patterns from the (c) x and (d) y polarization channels. Colorbar: phase (rad).

##### ii. Preparation of amyloid aggregates

We follow existing protocols [10, 11] for preparing amyloid fibrils. To aggregate the fibrils, we add 10 µM monomeric protein precursors (42 amino-acid residue amyloid-$\beta$ peptide) to an aggregation buffer of phosphate-buffered saline (PBS, pH 7.4), 150 mM NaCl, and 50 mM $Na_3PO_4$. We placed the mixture into an incubator at 37 °C with 200 rpm agitation for 24 h.

To image the amyloid fibrils, 10 µL the aggregated structures were placed into an ozone-cleaned cell culture chamber (Cellvis, C8-1.5H-N, No. 1.5H, 170 ± 5 µm thickness) for 1 h immediately after the incubation. After one hour, 200 µL of PBS solution containing 2.4 µM Nile Red (NR, Fisher Scientific, AC415711000) was placed into the amyloid-absorbed chambers for transient amyloid binding (TAB) [10] single-molecule orientation localization microscopy (SMOLM).

##### iii. Optical instrumentation and imaging procedure

The pixOL microscope is implemented using a home-built epifluorescence microscope as described previously [5, 11, 12]. Briefly, a polarization-resolved 4f imaging system, consisting of relay lenses (lenses 1-3) and a polarizing beamsplitter, is appended to a fluorescence microscope to project two polarized images onto separate regions of a camera (Fig. S11). A spatial light modulator (SLM, Meadowlark Optics, 256 XY Phase Series) is placed at the conjugate back focal plane (BFP) of the imaging system and loaded

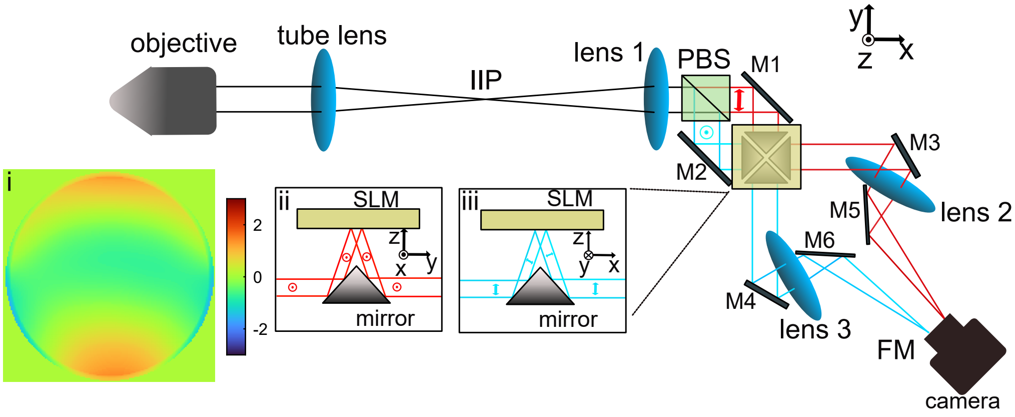

**Fig. S11.** Imaging system schematic. Single-molecule (SM) fluorescence is collected by the objective. A polarization-sensitive 4f system, comprising 3 lenses (lenses 1-3) and a polarizing beamsplitter (PBS), is added after the microscope's intermediate image plane (IIP). The PBS splits the light into x-polarized (red) and y-polarized (cyan) fluorescence. A pyramid mirror is used to reflect light from the two channels onto a spatial light modulator (SLM) placed at the conjugate back focal plane (BFP) of the microscope (insets ii and iii). The pixOL* phase mask (inset i) is loaded onto the SLM to modulate the phase of both channels simultaneously. Lenses 2 and 3 focus the x- and y- polarized fluorescence onto two non-overlapping regions of a single CMOS camera. Arrows denote the polarization of the light in each channel. M1-6, mirrors.

with the pixOL* phase mask (Fig. S11(i)) to modulate the x- and y-polarized fluorescence simultaneously (Fig. S11(ii,iii)).

The amyloid fibril is imaged with a 100× 1.4 NA oil-immersion objective lens (Olympus, UPlanSApo 100×). The samples were excited using a 561-nm circularly polarized laser (Coherent Sapphire, 1533 W/cm$^2$ peak intensity) tilted at ~30° from normal to excite both in-plane and out-of-plane oriented NRs. Fluorescence was collected by the same objective and filtered by a dichroic beamsplitter (Semrock, Di03-R488/561) and a bandpass filter (Semrock, FF01-523/610). Image stacks of 4,000 frames with 80 ms exposure were recorded.

##### iv. Two-channel image registration

Images simultaneously captured by the camera are required to registered before processing by Deep-SMOLM. The geometric transformation between the two channels was first roughly calibrated using fluorescent beads and then finely tuned using single-molecule (SM) imaging of amyloid fibrils. The fluorescent beads (Thermo Fisher Scientific, FluoSpheres, 0.1 µm, 580/605, F8801) are spin-coated on an ozone-cleaned coverslip (Marienfeld, No. 1.5H, 22 × 22 mm, 170 ± 5 µm thickness) and imaged using a polarization-sensitive standard DSF by turning off the pixOL* phase mask on the SLM shown in Fig. S11. We also captured 1,000 frames of single-molecule blinking on amyloid fibrils using the polarized standard DSF. The ThunderSTORM plugin [13] within ImageJ is used to localize the beads and single molecules within the two channels separately. The 2D bead positions across the two channels are then paired and used for calculating a global 2D polynomial transformation function using *images.geotrans.PolynomialTransformation2D* function in MATLAB. We then use the transformation function to pair single molecules on amyloid fibrils, and subsequently calculate a more accurate polynomial transformation function using the paired SM localizations.

To generate paired images for Deep-SMOLM estimation, we choose a field of view (FOV) within the y-polarized channel. For each pixel in the FOV in the y channel, we find the nearest corresponding pixel in the x channel using the transformation function and assemble these pixels together to form a transformed x-polarized image that is registered to the y-polarized image.

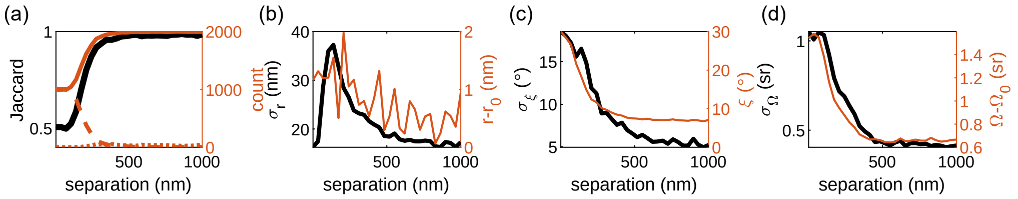

**Fig. S12.** Deep-SMOLM performance for estimating overlapping emitters with 500 detected signal photons and 2 background photons per pixel. (a) Deep-SMOLM (black) Jaccard index and the corresponding number of (orange solid) true-positive (TP), (orange dash) false-negative (FN), and (orange dot) false-positive (FP) emitters. (b) Deep-SMOLM (black) precision $\sigma_r$ and (orange) accuracy $r - r_0$ for estimating 2D position $\boldsymbol{r}$. (c) Deep-SMOLM (black) orientation precision $\sigma_\xi$ and (orange) absolute mean orientation bias $\xi$ (Eqn. 5). (d) Deep-SMOLM (black) precision $\sigma_\Omega$ and (orange) accuracy $\Omega - \Omega_0$ for measuring wobble angle $\Omega$.

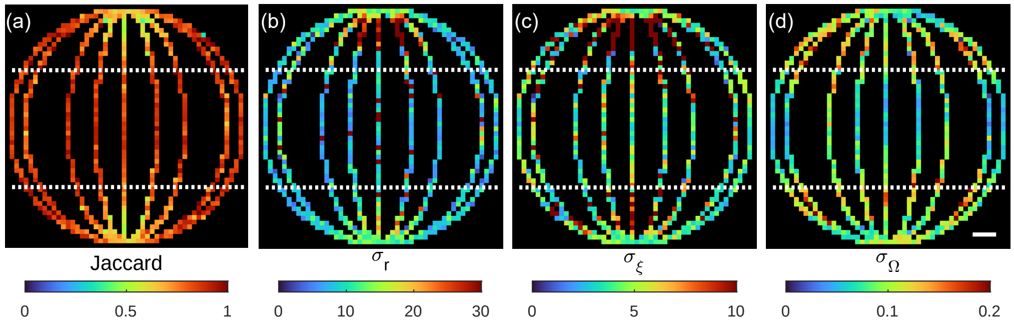

**Fig. S13.** Deep-SMOLM estimation performance for the simulated structure as quantified using (a) Jaccard index, (b) estimation precision $\sigma_r$ of 2D position, (c) estimation precision $\sigma_\xi$ of mean orientation angles $[\theta, \phi]$, and (d) estimation precision $\sigma_\Omega$ of wobble angle $\Omega$.

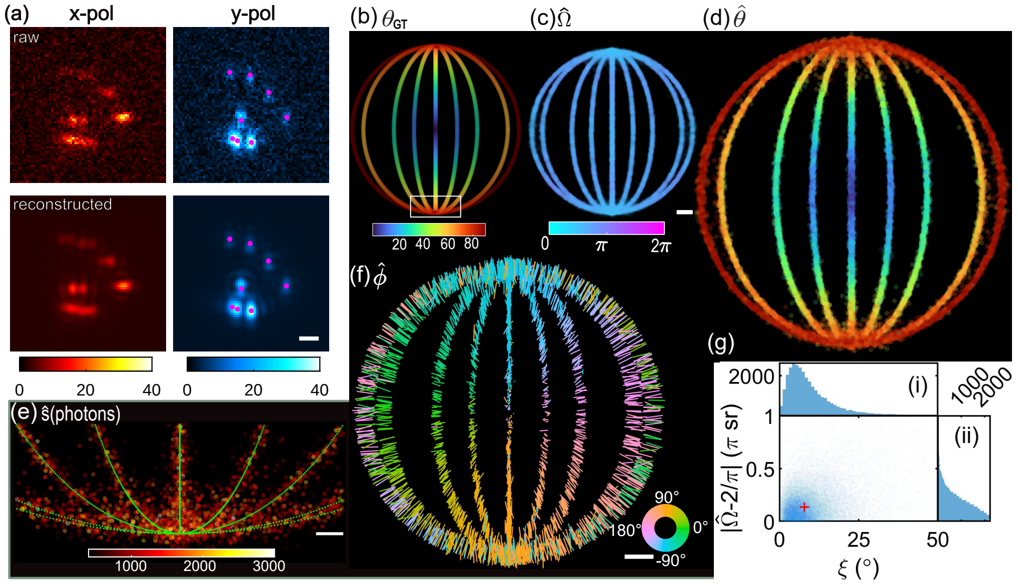

**Fig. S14.** Deep-SMOLM 5D imaging using simulated data with a 2 sr wobble angle. (a)(top) Simulated raw image compared to (bottom) images reconstructed from Deep-SMOLM estimates. Magenta dots: center position of each SM. (b) Synthetic structure containing nine 1D fibers color-coded with the ground truth polar angle $\theta_{\text{GT}}$. (c) Deep-SMOLM measured wobble angle $\hat{\Omega}$ (ground truth $\Omega_{\text{GT}} = 2$ sr). (d) Estimated polar angle $\hat{\theta}$. (e) Emitters within the white box shown in (b). Colormap: estimated signals $\hat{s}$ (photons). (f) Estimated azimuthal angle $\hat{\phi}$, where the length and direction of each line depict the magnitude of the in-plane orientation $\sin\hat{\theta}$ and direction of estimated azimuthal angle $\hat{\phi}$, respectively. The ground-truth orientations are perpendicular to the fibers. (g) Wobble angle estimation bias $|\hat{\Omega} - 2|$ versus mean orientation estimation bias $\xi$ (Eqn. 5). (Right) Distribution of wobble angle estimation bias and (top) mean orientation estimation bias. Scalebars: (c,f) 200 nm, (e) 50 nm.

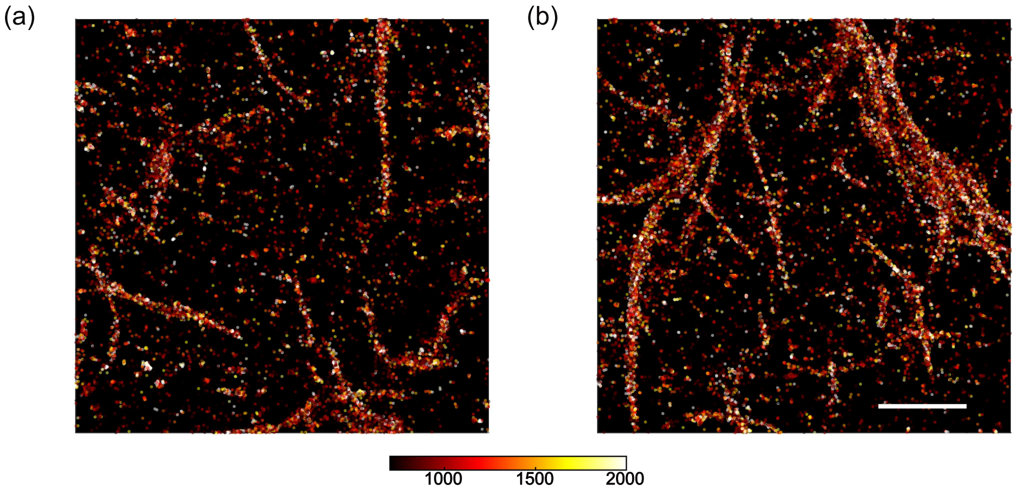

**Fig. S15.** Example SMLM reconstructions of amyloid fibrils. (a) A field of view with mostly separate amyloid fibrils. (b) A field of view with intertwined amyloid fibrils. Colorbars: signal photons for each detected emitter. Each SM is represented as a 2 nm filled circle. Scale bar: 1 µm.

Visualization 1 SM detection and position-orientation estimation using Deep-SMOLM for simulated biological fibers shown in Fig. 3. (Top) Simulated raw polarized images (red: x-polarized and blue: y-polarized) are compared to (bottom) images reconstructed using the 3D orientations and 2D positions estimated by Deep-SMOLM. Magenta dots: center position of each SM. Colorbar: photons/pixel. Scale bar: 1 μm.

Visualization 2 SM detection and position-orientation estimation using Deep-SMOLM for experimental amyloid fibrils shown in Fig. S15(a). (Top) Polarized images (red: x-polarized and blue: y-polarized) collected from the microscope are compared to (bottom) images reconstructed using the 3D orientations and 2D positions estimated by Deep-SMOLM. Colorbar: photons/pixel. Scale bar: 1 μm.

Visualization 3 SM detection and position-orientation estimation using Deep-SMOLM for experimental intertwined amyloid fibrils shown in Fig. S15(b). (Top) Polarized images (red: x-polarized and blue: y-polarized) collected from the microscope are compared to (bottom) images reconstructed using the 3D orientations and 2D positions estimated by Deep-SMOLM. Colorbar: photons/pixel. Scale bar: 1 μm.
